# Four-Dimensional Sparse Bayesian Tensor Decomposition for Gene Expression Data

**DOI:** 10.1101/2020.11.30.403907

**Authors:** Christopher C. Gill, Jonathan Marchini

## Abstract

Disease etiology may be better understood through the study of gene expression in four dimensional (4D) experiments that consist of measurements on multiple individuals, genes, tissues and under multiple conditions or through time. We have developed a sparse Bayesian four dimensional tensor decomposition method aimed at uncovering latent components or gene networks that could be linked to genetic variation. We used a Variational Bayes algorithm to fit the model which provides fast and accurate analysis. In this brief note we illustrate the utility of the method using simulated datasets, and show that when 4D data is available our method shows improved performance in estimating the true structure in the dataset, when compared to using a 3D method on a single slice of the 4D dataset. We also compare the results of the 4D method to that of the 3D method on a suitable unfolding of the dataset, demonstrating that similar performance is observed in this case, while the 4D method accurately recovers the additional structure in the data. We provide software that implements the method in R.

## Introduction

In the field of genomics the ever increasing size and availability of datasets is driven by technological advances and reductions in cost, and new techniques are required for their study. The availability of gene expression datasets in high dimensions is one such example, and single cell RNA-seq datasets have scale increasing exponentially [15]. Modern datasets can include multiple datatypes, providing different, complementary views of the samples and genome as described in [5]. The most basic gene expression dataset involves measurements in many individuals, samples, or cells, in a single tissue. This results in a two-dimensional dataset, with each datapoint indexed by an individual sample and a specific gene. Such datasets can be extended into three-dimensions by collecting data at different time points, or by integrating gene expression data from many tissues for each individual and gene. [5] point out that this additional dimension provides a different view of the underlying processes, in different contexts, so jointly analysing these together could provide more power to uncover signals in the data. For example, analysis of gene expression data has been used in [12] to understand the heterogeneity of expression in multiple tissues, and a longitudinal study of DNA methylation, [13], has been used to study the response to smoke exposure, identifying a critical window of in utero exposure.

It is notoriously difficult to perform inference on high-dimensional datasets, largely due to the challenges in handling the data, and finding a suitable model. One approach to making such datasets amenable to analysis is by way of fitting a latent underlying structure, explaining patterns of variability present in the data, including some model for the noise or confounding factors. This can include reducing the dimensionality of the problem, essentially finding a new representation of the data that captures the variability of the dataset with relatively few latent variables. Factor Analysis is one such approach, now widely used approach in genomics [14] and particularly in the case of a two dimensional gene expression dataset recorded as an *N* by *L* matrix *Y*. This results in a number, *C*, of vectors *x*^(*c*)^, of length *L* such that each individual’s gene expression data is modelled as a linear combination of the vectors *x*^(*c*)^. That is, each individual sample’s expression data is modelled as a set of individual weightings, or scores, for each component,

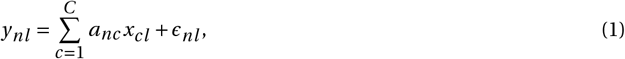

where *a_nc_* is the score for individual *n* and component *c*, *x_cl_* is the *l* th entry of *x*^(*c*)^, and *ϵ_nl_* is a Gaussian noise term. Thus, we describe this model as

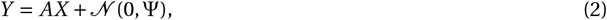

where Ψ is a diagonal covariance matrix. The rows of *X* (the vectors *x*^(*c*)^) are the latent factors, or gene loadings vectors, and the columns of the matrix *A* are the individual weightings, or individual scores vectors. In typical applications, the matrix A can be used for clustering, pseudo-temporal ordering, or further dimension reduction and visualisation. Typical dimension reductions for gene expression data are t-SNE, UMAP, or PCA [10, 11, 17]. The individual scores matrix can also be viewed as a ‘soft clustering’ that allows for continuous differentiation between samples, and from this viewpoint the individual scores matrix is often referred to as the pattern matrix [14]. The loadings matrix contains information pertaining to the sources of variation in the dataset across genes. Each component provides levels of expression of genes which are coexpressed across the dataset, and these can be further analysed for biological insights through enrichment analysis. For further discussion and references, see [14]. Unfortunately, even for two dimensional data sets, a basic factor analysis model lacks identifiability since any orthogonal change of basis preserves both the covariance and the decomposition. In practice, sparsity in the loadings matrix can facilitate rotational identifiability, and otherwise postprocessing techniques are used to identify rotations of the identified factors ([6]).

When considering a dataset of higher dimensions one can view it as a series of two dimensional datasets, for example as in Bayesian group factor analysis as described by [19] to model different data types or contexts while sharing a common individual scores matrix. This method has the disadvantages that the results may well reflect the choice of the two-dimensional slicing, each slice could be susceptible to the rotational identifiability issues mentioned above, and it introduces a large number of latent variables. An alternative approach that avoids this pitfall is to extend the factor analysis model to higher dimensions more directly. It can be shown that all tensors are sums of rank one tensors, and it is precisely such a decomposition which we focus on in the method we develop here. The concept of viewing a tensor as a sum of a finite number of rank one tensors appears in the literature in 1927, but only really became popular in 1970 when introduced as canonical decomposition (CANDECOMP) by [3], and Parallel Factor analysis (PARAFAC) by Harshman (see [6]). Research into methods aimed at decomposing data tensors to extract underlying explanatory structure has now been ongoing for at least four decades ([9]).

Consider a three-dimensional data tensor 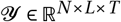, with measurements for *N* individuals, *T* tissues (or contexts, cell-types, time points etc.) and *L* genes. A PARAFAC decomposition of 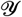 is described in equation 3 and illustrated in Figure 1,

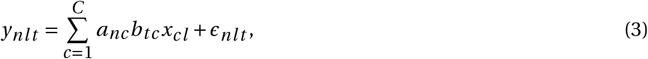

where *C* is the number of components, *A* ∈ ℝ^*N* × *C*^, *B* ∈ ℝ^*T* × *C*^, *X* ∈ ℝ^*C* × *L*^, and *ϵ_nlt_* is a noise term. This is typically summarised as a sum of rank one tensors, as

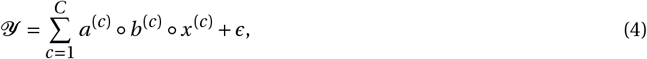

where ∘ is an outer tensor product, and *a*^(*c*)^, *b*^(*c*)^ are *c^th^* columns of *A*, *B* respectively and *x*^(*c*)^ is the *c^th^* row of *X*. A single component comprises all three of these vectors.

Another approach to three-dimensional tensor decomposition is the Tucker decomposition which includes an extra core tensor *P* ∈ ℝ^*C* × *C* × *C*^ where *C* is the number of components. This approach describes the tensor 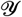 as a linear combination of tensors of the form *a*^(*i*)^ ∘ *b*^(*j*)^ ∘ *x*^(*k*)^, with coefficients given by the entries *p_i j k_* of *P*. The Tucker decomposition is more general than a PARAFAC decomposition (recovered by setting *p_i j k_* = 0 whenever *i*, *j*, *k* are not all equal, and 1 otherwise), but we note that the extra tensor *P* could introduce many variables that are not required in the PARAFAC decomposition, and may be less easily interpretable in a gene expression context.

**Figure 1:**
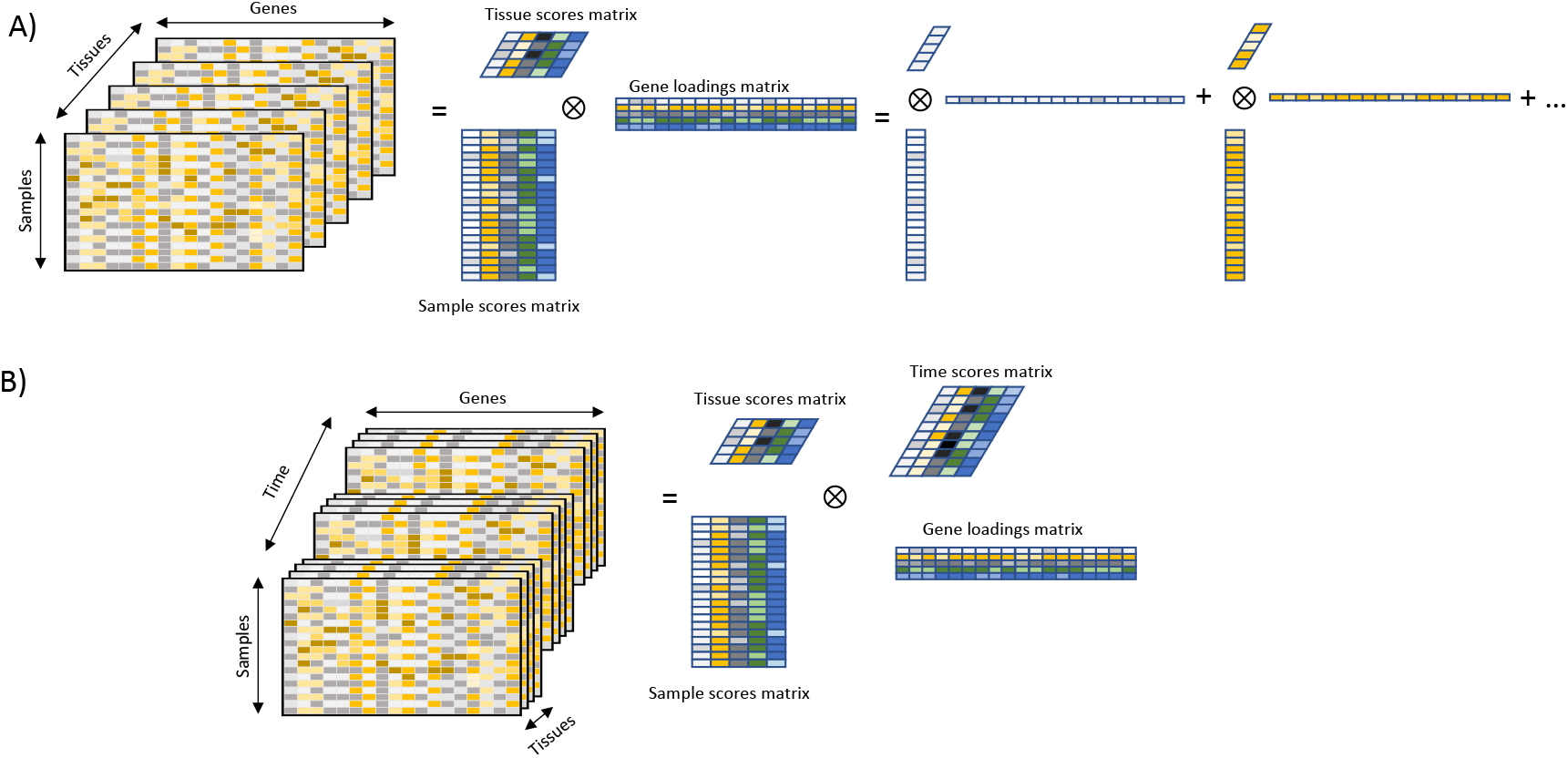
A) Diagram of a PARAFAC decomposition of a three dimensional tensor. B) Diagram of the four-dimensional PARAFAC model.

Recent work in [8] has developed a PARAFAC model for three-dimensional tensors of gene expression data. Under the assumption that biological processes involve relatively few genes, sparsity is induced in the gene loadings matrix *X* by means of a prior on the gene loadings incorporating a spike-and-slab distribution, which is a mixture of a point mass at 0, the ‘spike’, and a diffuse Gaussian centered at 0, the ‘slab’. Implemented using a fast Variational Bayes inference procedure, the resulting software was applied to the TwinsUK dataset and components were found which represented both meaningful biological sparse gene networks and experimental artefacts. Furthermore, genome-wide association studies of the component scores (columns of *A*) detected trans expression quantitative trait loci (eQTLs).

Although specifically designed for application to gene expression data, the model is very general and could be applicable to other contexts. Considering the increasing availability of high dimensional data in many fields, and in particular genetic datasets we have extended the three-dimensional parallel factor analysis model to four dimensions, maintaining the prior structure on the gene loadings matrix as developed in [8]. It is useful to consider this as time series data with a three-dimensional slice of the four-dimensional data tensor for each time point.

We describe the model in the Methods section, and the variational Bayes updates in the appendix. It is possible to restrict our four-dimensional implementation to recover the three-dimensional model from [8]. By application to simulated data, we show that a fourth dimension provides further information to decompose the latent structure in the tensor providing an accurate estimation of the latent structure. Comparison with results obtained from a 3D unfolding of the 4D tensors, shows that the three dimensional model can make effective use of the additional information to infer gene loadings and individual scores, but the main advantage of the 4D model is that it does infer the underlying latent structure in the data. The simulation is based on that from [8]. Software implementing our approach is available at https://github.com/marchinilab/SDA4D

## Methods

### The Model

The model is an extension of parallel factor analysis CANDECOMP/PARAFAC (CP) model to four dimensions with a sparse loadings matrix. This is a generalisation of the model used in [7] and [8] to four dimensions. Specifically, for a tensor 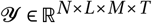, we consider the decomposition

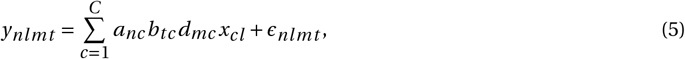

where *C* is the number of components, *A* ∈ ℝ^*N* × *C*^, *B* ∈ ℝ^*T* × *C*^, *D* ∈ ℝ^*M* × *C*^, *X* ∈ ℝ^*C* × *L*^ and *ϵ_nlmt_* is Gaussian noise. Figure 1 is an illustration of the model. The motivation is to consider a gene expression dataset of *N* individuals, *T* tissues, *M* time points, across *L* genes and we model the noise as having variance constant across individuals and time, with 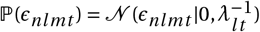, where *λ_l t_* is the precision, indexed by the tissue and gene.

Parallel factor tensor decompositions are invariant under permutation of the components. That is, if *P* is a permutation matrix and *A*, *B*, *D*, *X* are a PARAFAC decomposition of a tensor 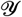, then *AP*, *BP*, *DP*, *PX* are also a PARAFAC decomposition of 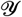. Furthermore, sign and scale are non-identifiable in a PARAFAC decomposition. We take into account these limitations when evaluating the results of the method in simulations. We follow the approach of [7] and with these limitations in mind, we use standard normal distributions for priors of *a_nc_*, *b_tc_*, *d_mc_*. We place a conjugate prior on *λ_l t_*, a Gamma distribution with parameters *u*, *v*.

Amongst a full spectrum of genes in any dataset, the number involved in any particular biological process will be small, so we aim to impose sparsity on the gene loadings vectors, using a ‘spike-and-slab’ prior, with a large density at zero, and remaining density spread as a normal distribution with mean zero and some precision. Assuming uniform precision across the component, the prior on *x_cl_* is defined as follows:

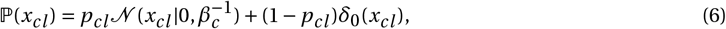

where *β_c_* is the precision of the ‘slab’, and *p_cl_* is a weight indicating whether the gene is active in that component. Following [16] we facilitate inference with such a distribution by expressing it as a product of a Gaussian distribution and a Bernoulli distribution. That is, we write *x_cl_* = *w_cl_s_cl_* where

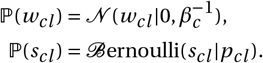

We apply a conjugate prior to *β_c_*, a Gamma distribution with parameters *e*, *f*. The mixing weight *p_cl_* is also given a spike and slab prior, this time involving a Beta distribution:

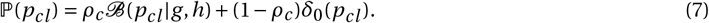

Such a hierarchical structure on the ‘spike’ was first introduced in [18], demonstrating that it reduced the false discovery rate in certain scenarios. Since *p_cl_* is the likelihood that gene *l* is ‘on’ in component *c* (that is, *x_cl_* ≠ 0), the value of *ρ_c_* determines the sparsity of the component. If *ρ_c_* takes a value close to 1 then *p_cl_* will take values from the Beta distribution, resulting in a dense component, whereas, if *ρ_c_* is close to zero then *p_cl_* has density highest at 0 and will result in a sparse component. We typically initialise with *g*, *h* close to zero resulting in a Beta distribution with peaks at zero and one, which further imposes sparsity. To make inference more straightforward, we write *p_cl_* = *ϕ_cl_ψ_cl_*, where

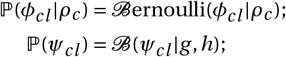

The prior for *ρ_c_* is a beta distribution with parameters *r*, *z*.

By setting *M* = 1, fixing 〈*d_mc_*〉 = 1 for all *c*, and neglecting to update *D*, we regain a three-dimensional model that is essentially equivalent to the method detailed in [8]. Since our method was implemented independently exact details may differ.

### Model Fitting

Due to the complex nature of this model, exact inference is not possible, so we have implemented a variational Bayes (VB), approximate inference, approach. In essence, VB approximates the posterior distribution by a distribution *Q* amongst a family by minimizing the Kullback–Liebler (KL) divergence between the posterior and the approximating distribution. In practice, this is achieved by maximizing a lower bound *F*(*Q*) to the model evidence, referred to as either the negative free energy, or the evidence lower bound (ELBO). A typical mean-field VB approach is to minimize the KL-divergence between the posterior and a fully factorized distribution, essentially imposing posterior independence on parameters. If the full family of parameters Θ is partitioned as Θ = ∪_*i*_*θ_i_* then *Q*(Θ) = Π*_i_ Q_i_* (*θ_i_*).

We take a coordinate ascent variational inference (CAVI) approach, implemented by initialising all *Qi*(*θi*) appropriately and iteratively updating each *Q_i_* to maximise *F*(*Q*) with respect to this term, while holding each of the approximating distributions fixed. When conjugate priors are used, the form for each optimal 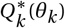 is analytical (has a recognisable closed form), and the distribution of 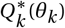 is dependent on the moments of *Q_i_*(*θ_i_*) for each *i* ≠ *k*. For parameters where conjugate priors are not used, and no analytical form of the distribution is available, point estimates for the parameters are used. In this case, standard optimisation techniques are used to maximise *F*(*Q*) over these variables (this is known as fixed form Variational Bayes).

The reader is referred to [2] and [1] for further details and references on Variational Bayes. We note that while Variational inference is known to be fast, and produce good posterior mean estimates, it typically underestimates the variance.

In our approach we follow [16] for inference with a spike and slab prior, and we find it beneficial to couple *w_cl_*, and *s_cl_* in the parameter partition as they together parameterize the spike and slab distribution.We partition the parameters as follows:

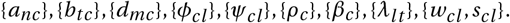

We further describe the resulting VB updates in the Appendix.

### A Simulation Study

We carried out a simulation study to illustrate the benefits of the 4D model, applied to 4D data, versus the 3D model applied to a single time point of a 4D dataset, and also against a 3D dataset derived from the 4D dataset.

We simulate 3D data for *N* = 200 individuals, across *L* = 500 genes, and *T* = 3 tissues with *C* = 8 underlying components. The data is generated by specifying matrices *A*, *B*, and *X* and adding noise. We generate an Individual Scores matrix *A* by sampling each entry from *N*(0, 1). *B* is the tissues matrix, and we give this a strong signal that differs strongly over tissues. Specifically, we simulate 8 components, across three tissues such that three components are active in just one tissue, three are active in two tissues and two are active in three tissues. The non-zero entries of *B* are sampled from {−1, 1}. The gene loadings matrix *X* is simulated by specifying a level of sparsity, *p*. Each entry of *X* is non-zero with probability *p*, in which case it is sampled from *N* (0, 1). Finally, each data point is generated as

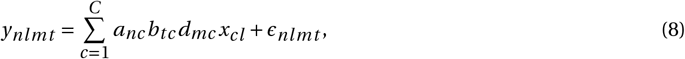

where *ϵ_nlmt_* is sampled from *N* (0, 10).

The 4D data was simulated in a similar manner, for *N* = 200 individuals across *L* = 500 genes, in *T* = 3 tissues, and across *M* = 16 time points. We simulated an underlying set of *C* = 8 components. *A*, *B* and *X* were generated as above, and we generated *D* deterministically, according to the following formulae (for *m* = 1,…, 16)

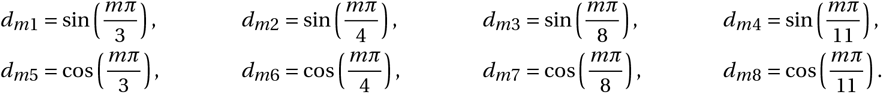

For each four dimensional tensor *Y*^(1)^ of dimension *N* × *L* × *M* × *T* we also unfold this into a three dimensional tensor *Y*^(2)^ of dimension *N* × *L* × *MT*. We consider the individuals and genes to be fixed, and the tissue and time dimensions to effectively capture context, which is how one might interpret the so-called tissue scores matrix in the three dimensional tensor. This results in an unfolding as 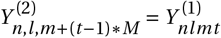. In this way, the unfolding determines a one-to-one correspondence between entries in the two tensors.

We varied the level of sparsity, such that *p* ∈ {0.1, 0.2, 0.3, 0.4, 0.5}, and at each level of sparsity we simulated 50 datasets. Variational inference is deterministic, and guaranteed to converge to a local maximum of the negative free energy, but can be sensitive to initialisation. For this reason we run the method ten times on each dataset and consider only the run which achieves the largest negative free energy at termination. For the purposes of the simulations we stop the algorithm as soon as the average rate of PIPs passing the threshold of 0.5 across 10-iterations drops below 1. The algorithm can shrink components to zero, so in the output, the algorithm selects the number of components. We initialised the code with double the true number of components in all cases, and fix the hyperparameters as follows: *e* = 10^−6^, *f* = 10^6^, *g* = 0, *h* = 0, *u* = 10^−6^, *v* = 10^6^, *r* = 1, *z* = 1 resulting in uninformative priors on precision of the noise and ‘slab’, a flat prior on the component sparsity parameter *ρ_c_*, and a beta with point masses at 0 and 1 for *ψ_cl_*.

The algorithm scales linearly in *N*, *L*, *M*, *T* and quadratically in the number of components *C*. Example running times are given in Table 1. Figure 2(A) shows the simulated tissue and temporal loadings, together with the estimates of these loadings when our algorithm was applied to a single simulated dataset, with sparsity level 0.3.

**Table 1:** Example running times. To demonstrate timing, we simulated two datasets with sparsity 0.4, simulated with 8 components each, and with dimensions (*N*, *L*, *M*, *T*) as given in the first column, identically to the simulated datasets used in the main text except extending the underlying tissue scores matrix *B* beyond *T* = 3 to *T* = 6 by drawing from a standard normal distribution. On a desktop computer running R version 3.5.2, we ran the method on each dataset twice, for 200 iterations, and to the convergence criterion laid out as used for the simulations in the main text.

| $(N, L, M, T, C)$ | Time for 200 iterations (s) | Time to convergence (s) |
| --- | --- | --- |
| (200, 500, 16, 3, 8) | 131 | 8 |
| (400, 1000, 32, 6, 16) | 1891 | 611 |

**Figure 2:**
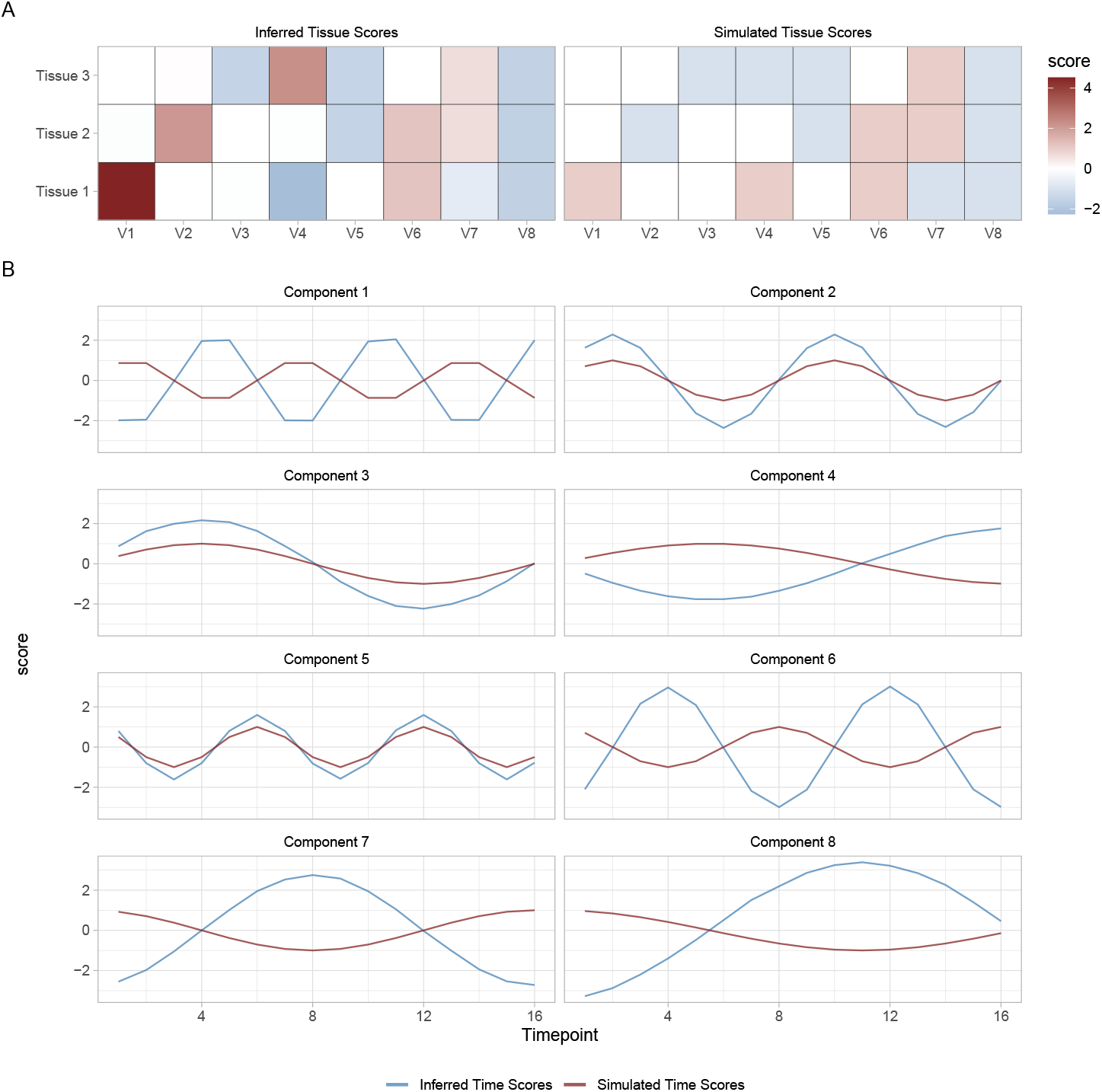
Example simulated and recovered tissue and time scores matrices. (A) The recovered and simulated tissue scores matrices. Component alignment was carried out with the individual scores matrix as described in the main text. Up to scale and sign (which are undetermined in the model), the tissue scores matrix is recovered well, and we note the scale is common on any given column reflecting the component-independence of scale in the model. (B) The corresponding recovered and simulated time scores for each component. Up to scale and sign these are recovered almost perfectly.

### Evaluating performance

We compared performance of the models by evaluating how well the method estimates the number of components and how well the individual scores and the gene loadings were recovered.

### Identifying estimated components

To evaluate the estimated number of non-zero components, we consider the posterior means 〈*s_cl_*〉, 〈*a_nc_*〉, 〈*b_tc_*〉, 〈*d_mc_*〉. The PIPs 〈*s_cl_*〉 are thresholded such that an entry 〈*s_c_l*〉 ≥ 0.5 indicates *x_cl_* ≠ 0 and gene *l* is active in component *c*. The posterior mean values 〈*s_cl_*〉 tend to be either close to zero, or to one, so although a threshold of 0.5 was used, any value between 0.1 and 0.9 gave very similar results.

Having identified the components that were shrunk to zero, those columns of *A*, *B*, *D* were removed. Computing the remaining postprocessing and metrics requires the correct number of components in all cases. If the number of estimated components was less than the true number, then additional zero components were added to *X*, and columns of zeros added to *A*, *B*, *D*. If more than 8 components were estimated then the components were selected to maximise the absolute correlation of the individual scores vectors with the true individual scores vectors.

The estimated individual, tissue, and time scores, and gene loadings matrices produced by this method are denoted *Â*, 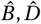, and 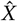 respectively.

### Individual scores vectors

Since there is a permutation and scale indeterminacy in the model we perform a search to determine the optimal permutation of components. That is, we permute the estimated components to maximise the average absolute correlation between the individual scores vectors, and change the sign of the vector in order to make the resulting correlations positive.

We compare the estimated individual scores vectors with the true individual scores vectors by root mean squared error. Since root mean squared error (9) is affected by scaling, each of the estimated and true individual scores vectors are scaled to have unit variance where the component is non-zero.

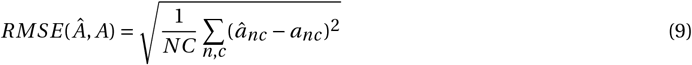

With a poor set of estimates, the optimal permutation may not be very well-defined (that is, it may not be significantly better than all others). In view of this, we use the sparse stability index as introduced in [4]. Let Σ be the *C* × *C* matrix with *i*, *j* entry the absolute correlation of the *i* th column of *Â* and the *j* th column of *A*. With this definition, the sparse stability index is defined as

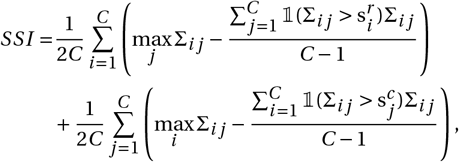

where s^*r*^, s^*c*^ are the vectors of row and column means of Σ respectively, and 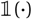 denotes an indicator function. This is invariant to scaling and permutation.

The SSI penalises cases where more than one entry in a row or column is close to one, or if a row or column contains no large entries. The former might happen if a component is split into two or more estimated components, and the latter if a component is missed. A high SSI indicates a good match between *A* and *Â*.

### Gene loadings vectors

Finally, we consider recovery of the set of non-zero gene loadings, by varying a threshold of inclusion on the posterior inclusion probabilities, calculating the false positive rate and true positive rate of the method and summarizing this as a receiver operating characteristic curve. The false positive rate (FPR) is calculated as in equation (10), and true positive rate (TPR) as in equation (11), where *X* is the simulated gene loadings matrix, and 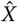 is the inferred gene loadings matrix, with columns aligned using the individual scores matrix. We define 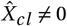 precisely when the posterior inclusion probability exceeds a defined threshold.

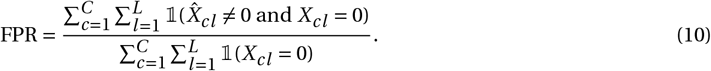

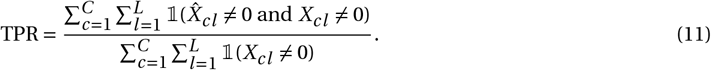

## Results

Figure 3 shows the results of the 3D and 4D models at estimating the number of components for each of the 5 levels of sparsity and each of the data types. We find that for sparsity levels *p* ≥ 0.2, the algorithm tends to overestimate the number of components in both 3D and 4D datasets, but this improves as *p* decreases (as the gene loadings matrix becomes increasingly sparse).

**Figure 3:**
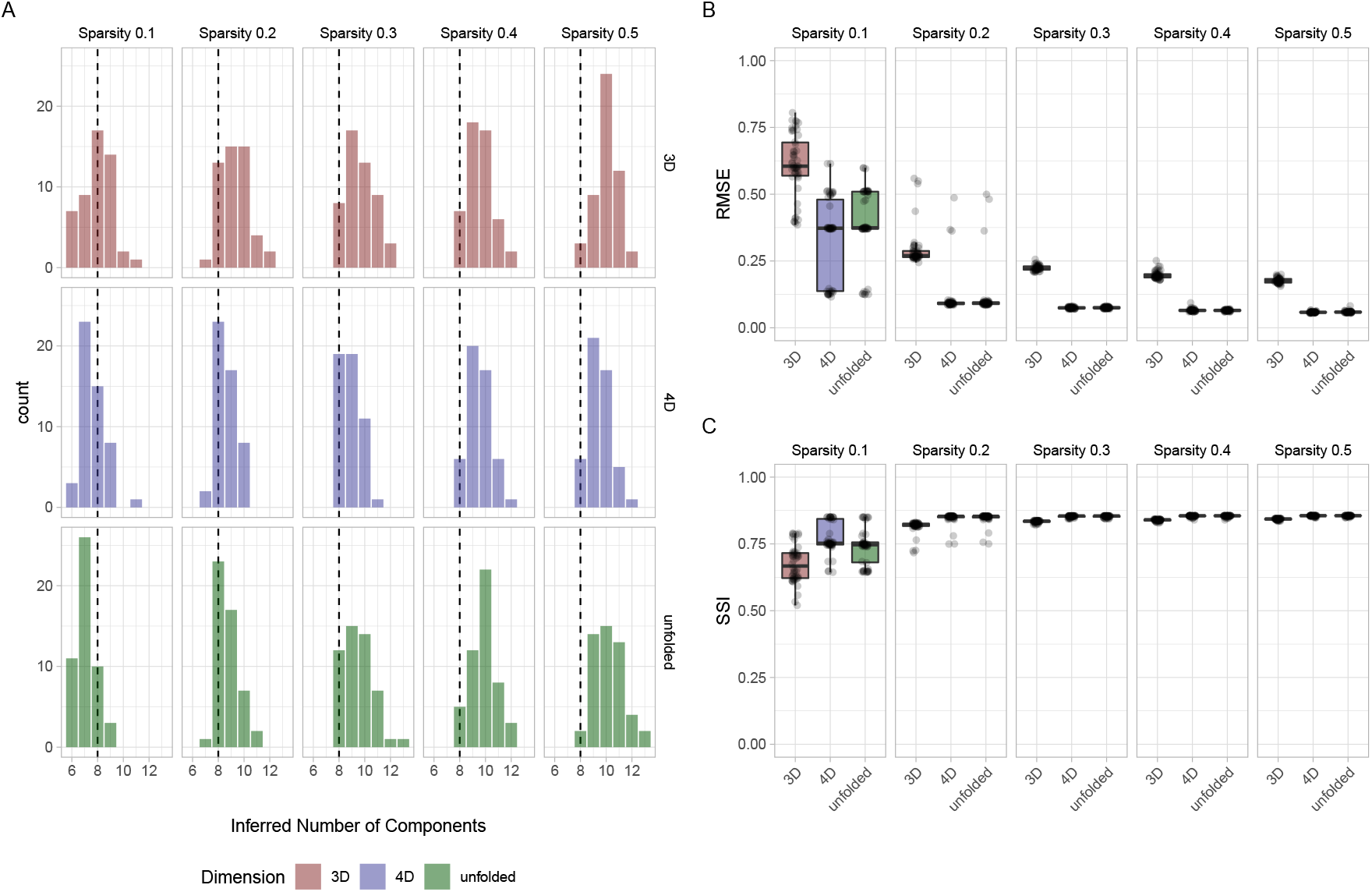
A) Comparison of estimated number of components. The dashed line indicates the true value of 8 underlying components. B) RMSE of the simulated and inferred cell scores after alignment as described in the main text. C) SSI of the inferred and simulated cell scores after alignment.

The number of components estimated in the 4D simulations typically showed less variability than the 3D simulation, and for each sparsity level, was typically closer to the true value. The 3D model on the unfolded dataset performs qualitatively similarly to the 4D model, with both typically overestimating the number of components for sparsity ≥ 0.2, when components are often split into two or more, and tending to underestimate the number of components for very sparse *p* = 0.1.

Figure 3(B,C) shows the RMSE and SSI metrics between estimated and simulated individual scores matrices for each sparsity level. We note that for 8 components, the SSI metric is bounded above by 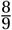. Both statistics appear stable and tightly distributed for sparsity between *p* = 0.2 and *p* = 0.5. It is noticeable that performance slightly worsens as the simulations become more sparse (low *p*), but the performance is particularly less predictable at *p* = 0.1. The RMSE achieved by the four-dimensional algorithm is generally lower than that achieved by the three-dimensional algorithm and the SSI is generally higher in the four dimensional results than those of the three-dimensional results, highlighting the improved performance achieved by incorporating time-series data. However, the 3D model applied to the unfolded datasets perform very similarly to the 4D model in all but the sparsity 0.1 model, where the 4*D* model slightly outperforms the unfolded results.

We note also that in order to calculate the statistics we use here, we have discarded estimated components since the algorithm often inferred more than the 8 simulated components. It is perfectly possible for this method to split a component into two or more inferred components. This would not be detected in the statistics used here and could lead to decreased sparse stability index, or increased root mean squared error.

We also compare the false-positive-rate and power achieved by each method across sparsity levels in Figure 4. In all cases, the false positive rate is very conservative. These results show a similar pattern to the other statistics, a decrease in performance as sparsity increases. The four-dimensional algorithm again seems comparable to the three-dimensional results from the unfolded data, and clearly outperforms the three-dimensional algorithm on a smaller dataset, with increased true positive rate and lower false positive rate across the majority of simulations (in fact all simulations for *p* = 0.3, 0.4, and *p* = 0.5).

**Figure 4:**
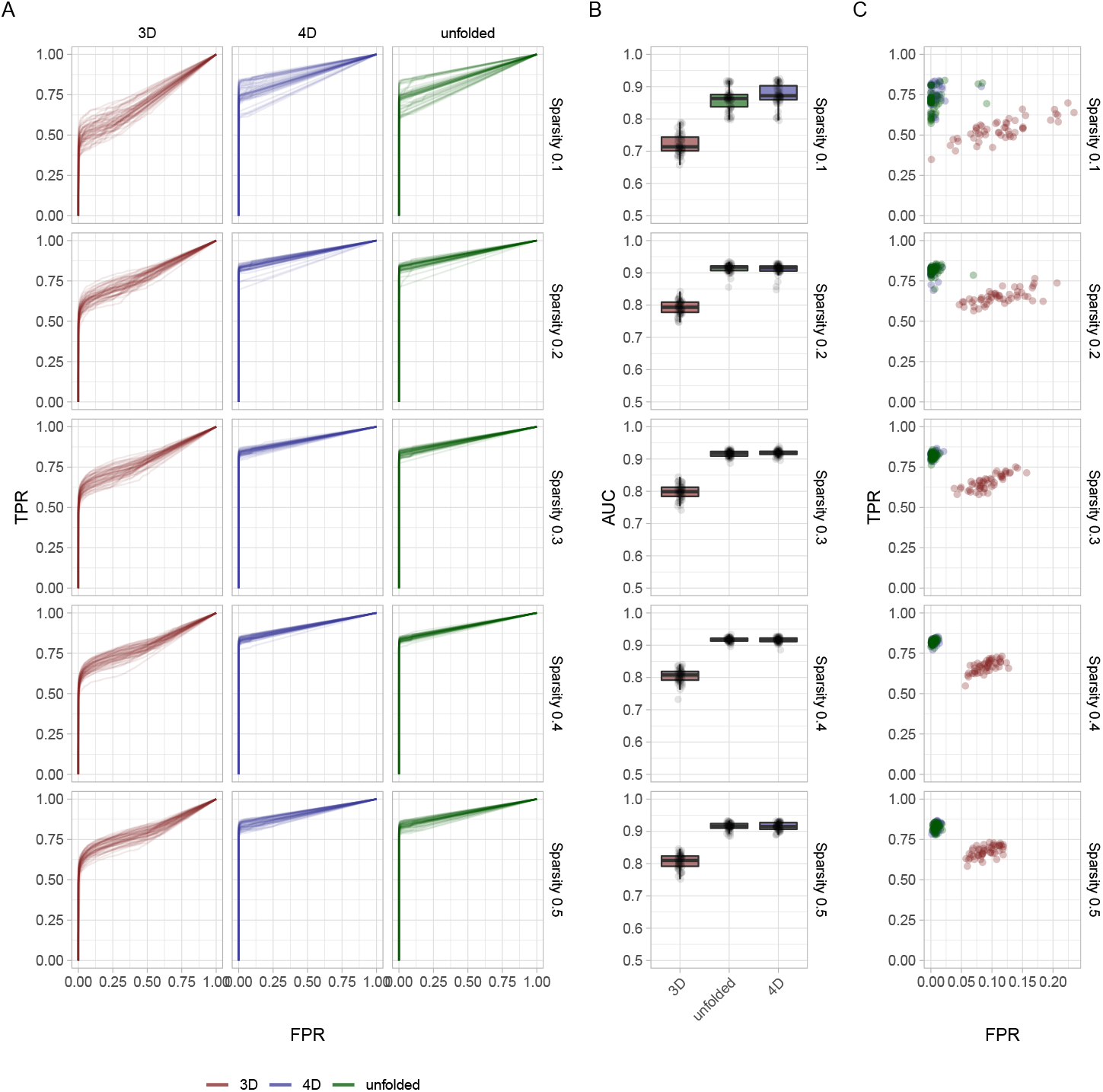
Comparison of recovery of the gene loadings matrix by FPR and TPR. On the 50 datasets at each level of sparsity we consider false positive rate and true positive rate for both 3*D* and 4*D* simulations. A) ROC curves for the three methods for the highest ELBO run on each dataset/method after component alignment using individual scores as described in the main text. B) Area under the curve for the ROC curves. C) FPR/TPR plotted when applying a threshold of 0.5 to the posterior inclusion probabilities.

We note also that the time and tissue scores matrices have been recovered very well in all scenarios. An example of the tissue scores matrix as simulated and as recovered is shown in Figure 2(A). For the same dataset and run, we plot the recovered time scores in Figure 2(B). In both plots the scale and sign indeterminacy of the model is clear, but subject to this consideration, the recovery is excellent. These components were aligned with the simulated components by the individual scores matrix, following the method described earlier in the text. Applying this same method and recording the average absolute correlation achieved across the columns of the estimated and simulated time scores matrices, we found exceptionally clear recovery of the time scores matrix (Figure 5). We consider this to be a relatively complex time scores matrix, with several columns highly correlated, but it was recovered extremely well.

**Figure 5:**
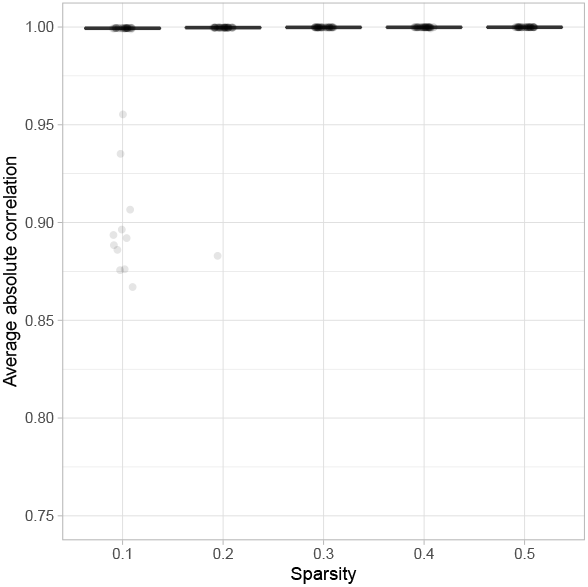
Time Scores matrix is recovered well. For the highest negative free energy runs on each dataset we plot the average absolute correlation of the non-constant columns of the estimated time scores matrix with the true underlying time scores matrix, based on the component alignment carried out with the Individual scores matrix described in the main text. The plot above shows excellent recovery of the matrix. We note also that the sign and scale are indeterminate in the model, as shown in Figure 2.

## Discussion

The application of sparsity inducing priors to a parallel factor analysis model for gene expression data was novel in and has been shown to have application in large gene expression studies, in particular having power to detect trans eQTLs. We have extended this model to perform inference on four-dimensional data with gene expression data in mind, although we expect the approach to be more widely applicable.

The interpretation of the matrix *B* has been that of a tissue scores matrix and this has been recovered well. Similarly the matrix *D*, which we have used to define the variability of component expression over time has been recovered well - see for an example Figure 2 and Figure 5. The combination of these two matrices and patterns being recovered is indicative of the power of both methods to detect ‘discrete’ patterns in the componentwise scores such as that seen in the tissue scores matrix, as well as the more continuous variation as captured in the time scores matrix.

We have shown that the four-dimensional method outperforms the three-dimensional method in recovering the gene loadings in components and the individual scores matrix, and observed excellent recovery of the tissue and time scores matrices. The results presented here suggest that the results from the 3D model are limited by the amount of data in a single time slice. When applied to a suitable unfolding of the four dimensional data, the results of the 3D model are comparable with that of the 4D model in gene loadings and individual scores. However, the 4D parafac method presented here reliably recovered the tissue and time scores across multiple components and simulations.

While we present here a brief summary of results from a simulation study similar to that of [8], we have observed qualitatively identical results in another scenario designed to be more symmetric in dimension, with a range of dimensions up to 50 × 50 × 50 × 50 datasets.

We expect that with the increasing availability of gene expression data, time series data across multiple tissues will become available and this method will prove invaluable in decomposing such datasets into active components with weightings (scores) for different features (dimensions). This approach has the potential to provide insight into components of genes involved in biological processes, as well as their differential expression over time.

## Acknowledgements

We are grateful to Victoria Hore for useful conversations about the SDA method and software implementation. CG also gratefully acknowledges support from the EPSRC Doctoral Training Grant EP/G037280/1.

## Appendix

We summarise the model as follows for a data tensor 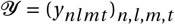:

## The Likelihood

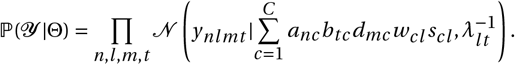

## Prior distributions

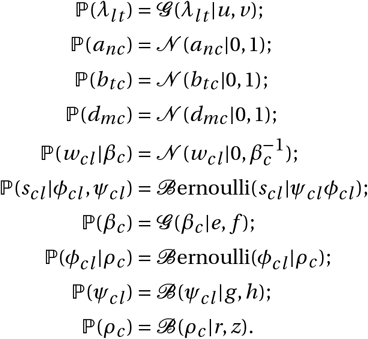

In deriving our Variational inference algorithm, we partition the variables as

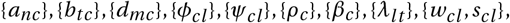

so assume all variables are independent except *w_cl_*, *s_cl_* which remain coupled. This results in the following VB updates, where we denote 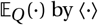:

## Individual scores matrix

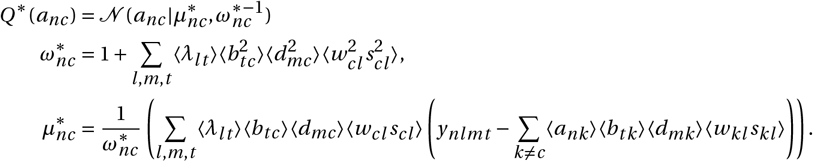

## Tissue scores matrix

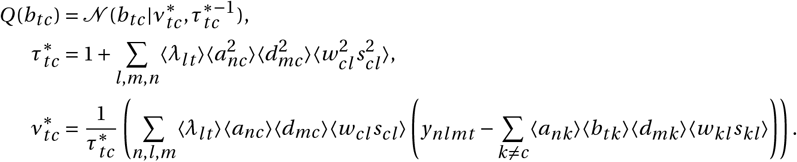

## Time scores matrix

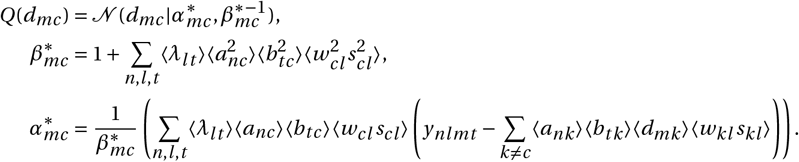

## Noise precision

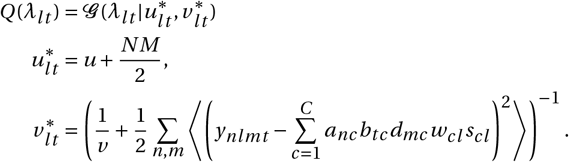

## Gene loadings matrix

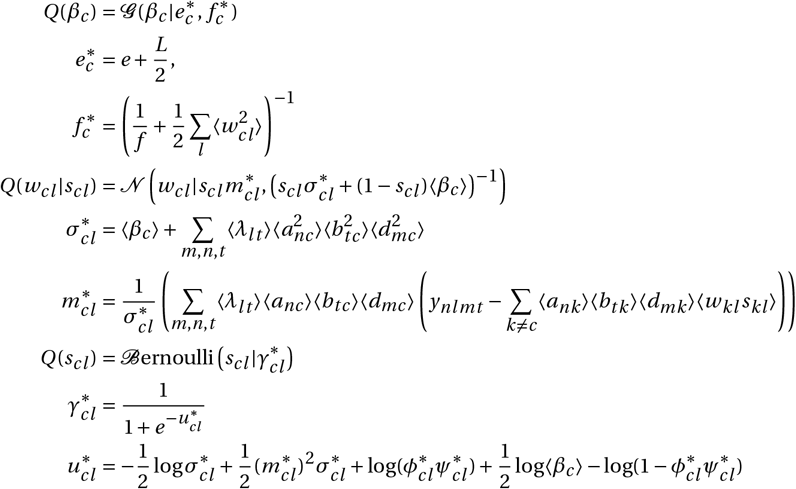

Following the same procedure for the variables *ρ_c_*, *ψ_cl_*, *ϕ_cl_* does not result in a closed form distribution. In this case, we maximise the ELBO with respect to point estimates of these parameters. The corresponding component of *F*(*Q*), dependent on these parameters, is 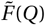 defined as follows:

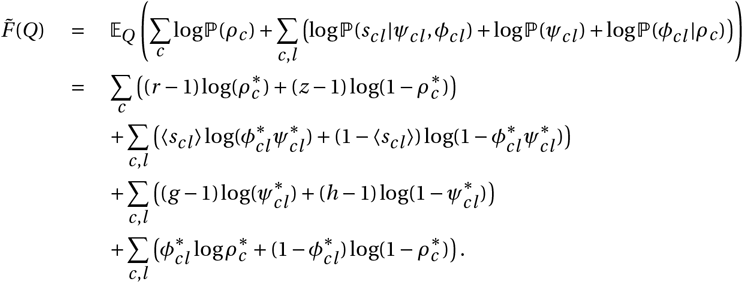

There is a unique solution to 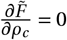, which gives the update for *ρ_c_* as

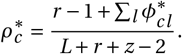

For the remaining parameters *ϕ_cl_*, *ψ_cl_* we optimise *F*(*Q*) by maximising 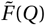 using Newton’s method in two dimensions. The gradient vector is

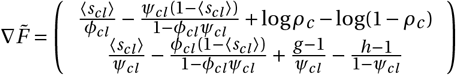

The Hessian matrix, *H*, is calculated as

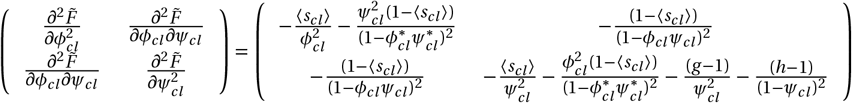

The update for (*ϕ_cl_*,*ψ_cl_*) is then

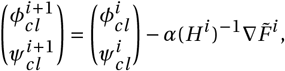

where *α* is chosen by a backtracking line search to ensure the step increases 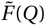.

Each of these updates is guaranteed to increase the ELBO,

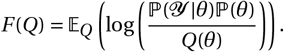

The ELBO (or negative free energy) for this model is as follows:

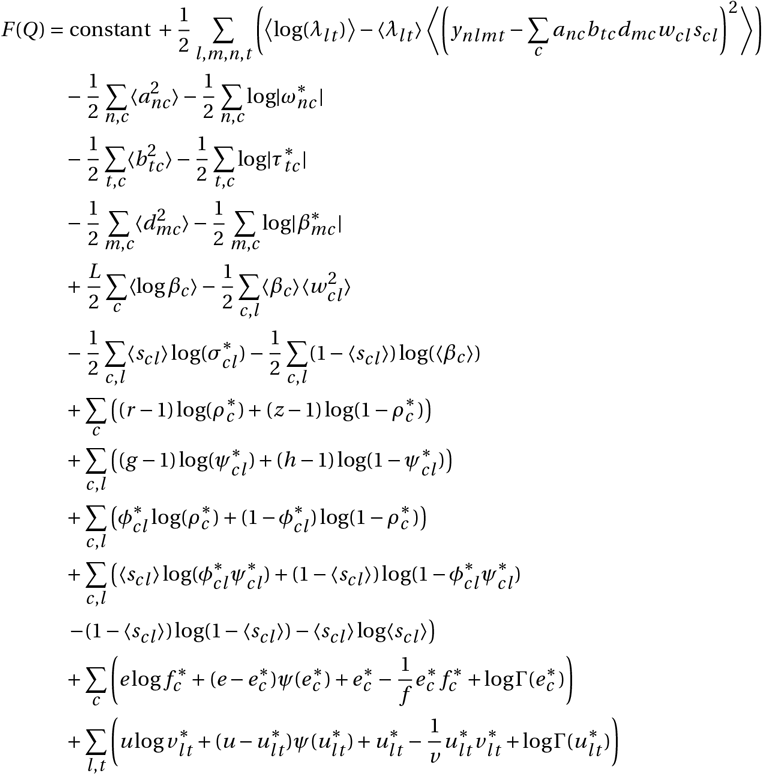

Here *ψ* is the digamma function.

